# A complex regulatory cascade involving quorum sensing regulates siderophore-mediated iron homeostasis in *Chromobacterium violaceum*

**DOI:** 10.1101/2023.12.13.571441

**Authors:** Bianca B. Batista, Vinicius M. de Lima, Beatriz A. Picinato, Tie Koide, José F. da Silva Neto

## Abstract

Iron is a transition metal used as a cofactor in many biochemical reactions. In bacteria, iron homeostasis involves Fur-mediated de-repression of iron uptake systems, such as the iron-chelating compounds siderophores. In this work, we identified and characterized novel regulatory systems that control siderophores in the environmental opportunistic pathogen *Chromobacterium violaceum*. Screening of a 10,000-transposon mutant library for siderophore halos on PSA-CAS plates identified seven possible regulatory systems involved in siderophore-mediated iron homeostasis in *C. violaceum*. Further characterization revealed a regulatory cascade that controls siderophores involving the transcription factor VitR acting upstream of the quorum sensing (QS) system CviIR. Mutation of the regulator VitR led to an increase in siderophore halos, and a decrease in biofilm, violacein, and protease production. We determined that these effects occurred due to VitR-dependent de-repression of *vioS*. Increased VioS leads to direct inhibition of the CviR regulator by protein-protein interaction. Indeed, insertion mutations in *cviR* and null mutation of *cviI* and *cviR* led to an increase of siderophore halos. RNA-seq of the *cviI* and *cviR* mutants revealed that CviR regulates CviI-dependent and CviI-independent regulons. Classical QS-dependent processes (violacein, proteases, and antibiotics) were activated at high cell density by both CviI and CviR. However, genes related to iron homeostasis and many other processes were regulated by CviR but not CviI, suggesting that CviR acts without its canonical CviI autoinducer. Our data revealed a complex regulatory cascade involving QS that controls siderophore-mediated iron homeostasis in *C. violaceum*.

**Importance:** The iron-chelating compounds siderophores play a major role in bacterial iron acquisition. Here, we employed a genetic screen to identify novel siderophore regulatory systems in *Chromobacterium violaceum*, an opportunistic human pathogen. Many mutants with increased siderophore halos had transposon insertions in genes encoding transcription factors, including a novel regulator called VitR, and CviR, the regulator of the quorum sensing (QS) system CviIR. We found that VitR is upstream in the pathway and acts as a dedicated repressor of *vioS*, which encodes a direct CviR-inhibitory protein. Indeed, all QS-related phenotypes of a *vitR* mutant were rescued in a *vitRvioS* mutant. At high cell density, CviIR activated classical QS-dependent processes (violacein, proteases, and antibiotics production). However, genes related to iron homeostasis and type-III and type-VI secretion systems were regulated by CviR in a CviI- or cell density-independent manner. Our data unveil a complex regulatory cascade integrating QS and siderophores in *C. violaceum*.

## Introduction

Iron is an essential micronutrient required by almost all living organisms since it acts as a cofactor for enzymes involved in crucial biological processes. ^1,2^ Bacteria uptake iron from different sources and in distinct ways.^3^ While Fe^2+^ is directly transported by systems located in the cytoplasmic membrane, such as FeoAB and EfeUOB, the insoluble form Fe^3+^ is solubilized and transported as siderophore-Fe^3+^ complexes. Siderophores are low molecular weight molecules with high affinity for Fe^3+^.^4,5,6^ In Gram-negative bacteria, Fe^3+^-siderophore complexes are transported across the outer membrane by TonB-dependent receptors, and from the periplasm to the cytoplasm by ABC-type transporters.^2,7^

Bacteria maintain iron homeostasis by regulating gene expression in response to iron availability. In most bacteria, this is an orchestrated mechanism involving Fur, an iron-sensing global transcription factor, and iron-responsive small regulatory RNAs (sRNA).^8,9,10^ When a sufficient amount of iron is present, the Fur-Fe^2+^ metalloprotein complex represses genes encoding iron uptake systems by binding to a specific DNA sequence known as Fur box, located in their promoter regions.^1,8,10,11^

Other regulatory mechanisms controlling iron homeostasis have been identified. In *Xanthomonas campestris*, a virulence-associated global regulator called XibR positively regulates motility and iron uptake and storage, while it negatively regulates siderophore synthesis in response to iron levels.^12^ Iron homeostasis can be integrated into quorum sensing (QS) circuits, a cell-cell communication process in which cells produce, detect, and respond to signaling molecules called autoinducers.^13^ Considering that siderophores are known as public goods, it is not surprising that the QS systems of some bacteria regulate siderophore production.^14,15,16,17^

*Chromobacterium violaceum* is a Gram-negative, saprophytic bacterium found in the soil and water of tropical and subtropical regions;^18^ it is an opportunistic pathogen that causes severe infections in humans.^19,20^ *C. violaceum* produces a violet-colored pigment called violacein, which has been shown to have antibacterial, antiparasitic, antiviral, and antitumor actions *in vitro*.^21,22^ In *C. violaceum*, the violacein production is activated by the CviIR QS system, in which the CviI enzyme produces N-acyl-L-homoserine lactone (AHL) autoinducers.^23^ At high cell density (HCD), the AHLs accumulate and bind to the CviR regulator, which in turn regulates several processes.^24,25,26,27^ A CviR DNA binding site was mapped upstream of the violacein biosynthesis operon and used for *in silico* prediction of other potential CviR-regulated genes.^25^ However, the global CviR regulon and the connection between QS and iron homeostasis remains unexplored in *C. violaceum*.

Our group has shown that *C. violaceum* synthesizes at least two catecholate siderophores (chromobactin and viobactin) required for *C. violaceum* virulence.^28^ We propose that these siderophores are assembled by the nonribosomal peptide synthetases (NRPS) CbaF and VbaF from the 2,3-DHBA precursor and imported by the TonB-dependent receptors CbuA and VbuA, respectively.^28^ In another study, we have demonstrated that *C. violaceum* uses heme via the ChuPRSRTUV system, and that both siderophores and heme are important iron acquisition strategies during infection.^29^ Further work by our group has shown that Fur protects *C. violaceum* against iron overload and oxidative stress. Also, Fur represses genes related to iron homeostasis and controls virulence in this bacterium.^30^ However, it remains unknown whether other transcription factors regulate the production and uptake of siderophores in *C. violaceum*.

In this work, we identified novel regulatory mechanisms involved in iron homeostasis in *C. violaceum* by screening a transposon mutant library for altered siderophore halos in PSA-CAS plates. Our data unveil a regulatory cascade involving the transcription factor VitR that culminates in the QS system CviIR controlling siderophore-mediated iron homeostasis.

## Results

### A global transposon mutagenesis approach reveals novel regulatory systems involved in siderophore-mediated iron homeostasis in *C. violaceum*

Fur, a master iron-responsive regulator, represses siderophore production and utilization in *C. violaceum*.^30^ To identify novel regulatory systems controlling siderophores, we used the T8 transposon^30,31^ to generate a library of 10,000 transposon mutants in *C. violaceum* ATCC 12472. Library screening on siderophore-indicative PSA-CAS plates revealed 132 transposon-mutant strains with altered siderophore halos (101 strains with increased halos and 31 strains with decreased ones) (Table S1). Sequencing of semi-degenerate PCR products from the 132 mutant strains identified unique transposon insertion sites in 25 different genes in the *C. violaceum* genome, with some genes showing multiple independent transposon insertions (Table S1). Mutated genes grouped into different functional categories, and six encoded regulatory systems (Figure 1A; Table S1). We focused on three of these regulatory systems: (I) the transcription factor VitR (CV_1057) (Figure 1B); (ii) the two-component system AirSR (CV_0536-37) (Figure 1C); and (III) the transcription factor CviR (CV_4090) of the QS system CviIR (Figure 1D). For all these genes, we further confirmed the presence of transposon insertions by PCR, and we generated null mutant strains (Figure 1B-D). These results indicate that several regulatory systems control siderophore-mediated iron homeostasis in *C. violaceum*.

**Figure 1.**
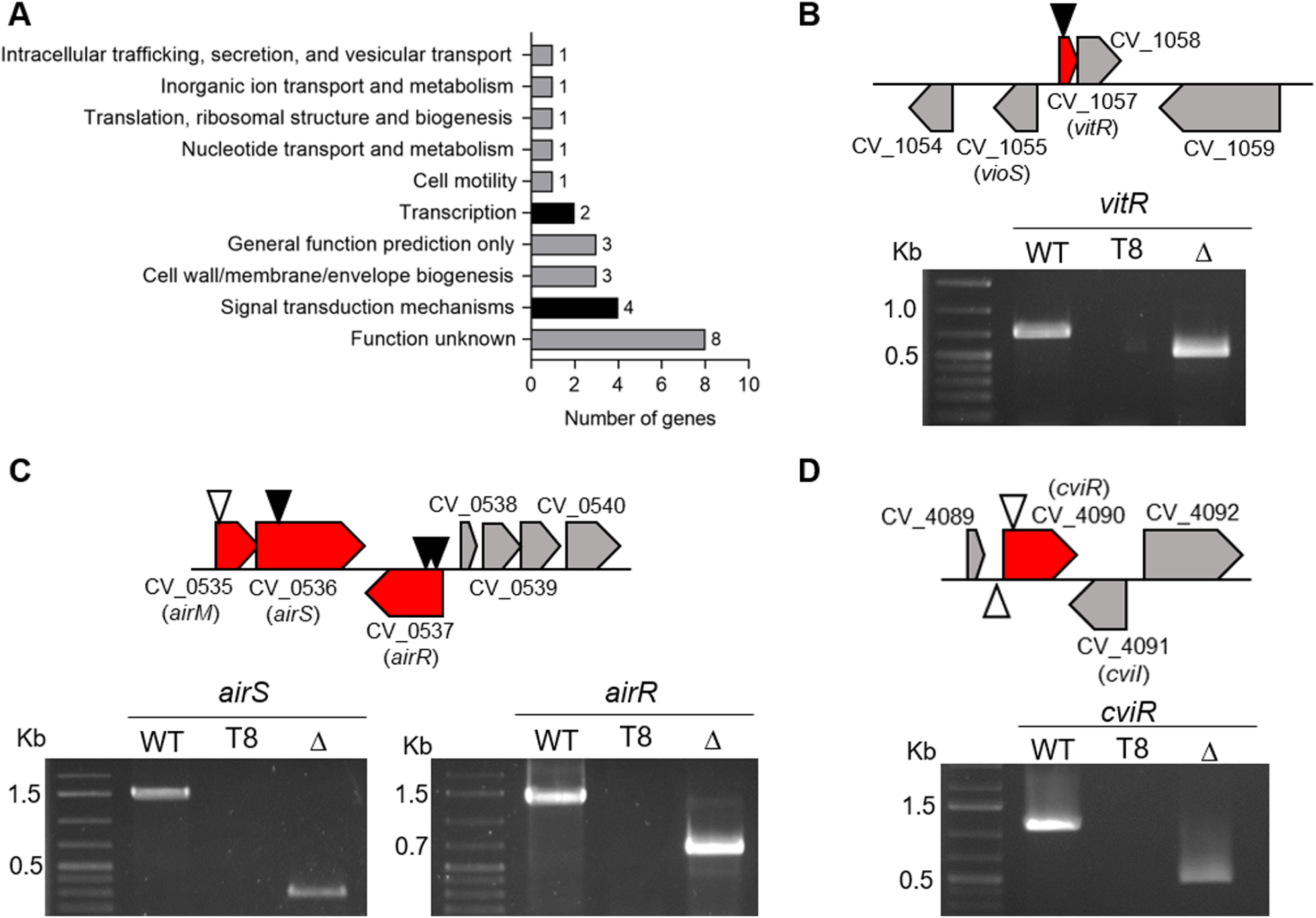
Functional classification of the genes with transposon insertion and identification of the insertion site in mutant strains of regulatory genes. **A.** Functional classification of 25 genes with transposon insertion that affected siderophore activity. **B-D.** Gene organization of the regulatory systems with transposon insertions, and PCR to confirm the insertion and the null-mutant strains. Black arrowhead, one insertion site; white arrowhead, multiple insertion sites; WT, wild-type strain; T8, transposon mutant strain of the indicated gene; Δ, null-mutant strain of the indicated gene; molecular weight marker 1 Kb plus DNA Ladder (Thermo Scientific).

### The two-component system AirSR plays a role in siderophore homeostasis

Transposon insertions into the genes CV_0535, CV_0536, and CV_0537 increased the siderophore halos (Table S1; Figure 1C). Recently, the orthologs of CV_0535-36-37 in *C. violaceum* ATCC 31532 were characterized as an antibiotic-induced response system (Air system) composed of an oxidoreductase (AirM), a histidine kinase (AirS), and a response regulator (AirR). The Air system acts via the CviIR signaling pathway to activate violacein production.^32^ In agreement with our findings on the transposon-mutant strains, the null-mutant strains Δ*airS*, Δ*airR*, and Δ*airSR* showed an increase in siderophore halos; these phenotypes were reversed by complementation (Figure 2A and B). These results indicate that the two-component system AirSR controls siderophore homeostasis in *C. violaceum*.

**Figure 2.**
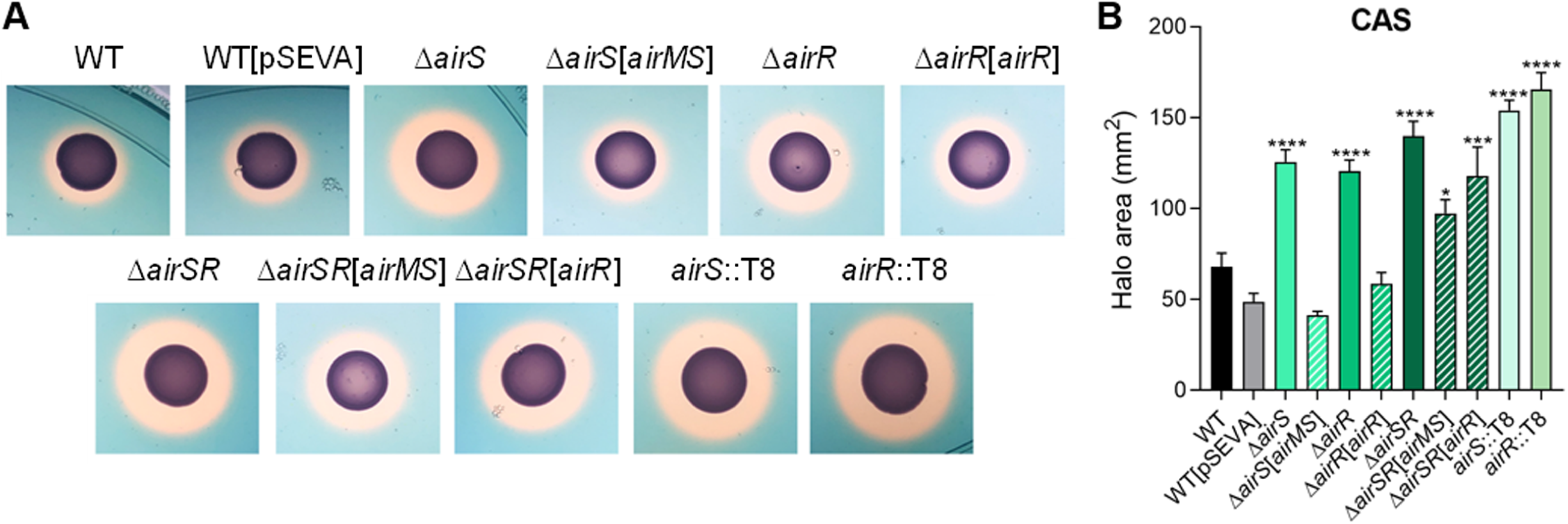
Characterization of the two-component system *airSR*. **A.** CAS (Chrome Azurol S assay for siderophores) tests showing the production of siderophores (orange halos) by the indicated strains inoculated on PSA-CAS plates. We generated null mutant strains for the *airSR* system with the same phenotype as the *airS*::T8 and *airR*::T8 strains. **B.** Measurement of the CAS halo areas of the indicated strains. Data from three biological assays. Statistical analyses using One-way ANOVA followed by Holm-Sidak’s multiple comparisons test. * p<0.05; ** p<0.01, *** p<0.001, **** p<0.0001, when not indicated, n.s. (not significant).

### The transcription factor VitR controls siderophore, violacein, and biofilm formation in *C. violaceum*

Transposon insertion in CV_1057 resulted in increased siderophore halos (Table S1; Figure 1B). The gene CV_1057 encodes a putative transcription factor belonging to the superfamily Cro, family XRE, that we named VitR (<u>v</u>iolacein inhibitor <u>r</u>egulator). A Δ*vitR* mutant strain showed increased siderophore halos, validating the phenotype of the transposon mutant (Figure 3A and B). Interestingly, we observed that when grown in LB broth for 24 hours, the Δ*vitR* mutant produced less violacein than the wild-type (WT) strain (Figure 3C). Also, Δ*vitR* formed less biofilm than the WT strain (Figure 3D). Growth curves indicate that Δ*vitR* had the same growth in LB (Figure 3E), but a slight growth decrease in LB-chelated for iron (150 µM DP) (Figure 3F), when compared to the WT strain. All observed phenotypes were rescued in a Δ*vitR*-complemented strain (Figure 3). Together, these data indicate that VitR regulates siderophores, violacein production, and biofilm formation in *C. violaceum*.

**Figure 3.**
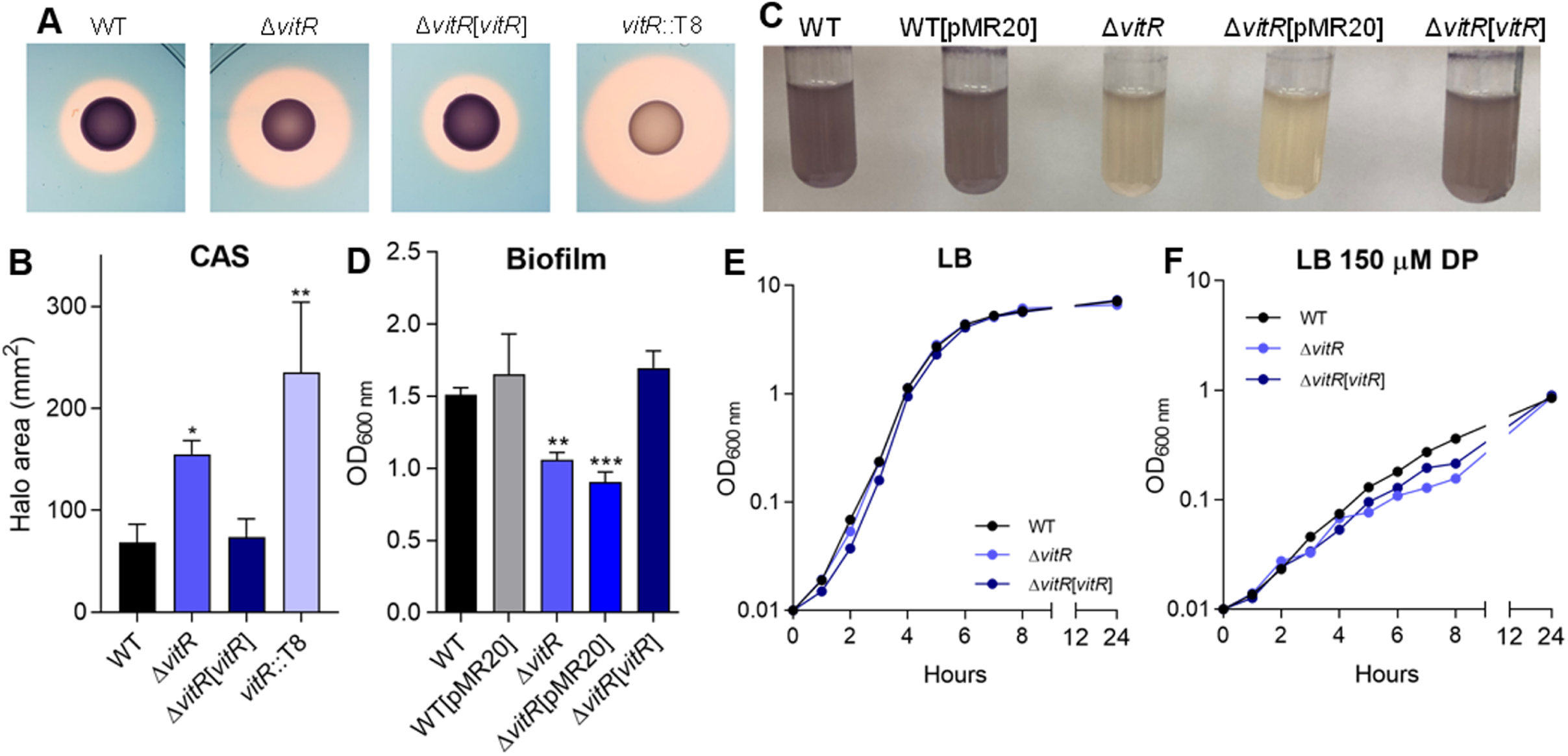
Phenotypic characterization of the *vitR* regulator. **A.** CAS tests showing the production of siderophores (orange halos) by the indicated strains inoculated on PSA-CAS plates. We generated a null mutant strain for the CV_1057 (*vitR*) gene with a similar phenotype as the *vitR*::T8. **B.** Measurement of the CAS halo areas of the indicated strains. Data from three biological assays. Statistical analyses using One-way ANOVA followed by Dunnett’s multiple comparisons test. * p<0.05; ** p<0.01, when not indicated, n.s. (not significant). **C.** Growth of the indicated strains in LB medium to verify the production of violacein. **D.** Biofilm assay of the indicated strains. The strains were grown in LB medium for 24 hours and the assay was performed with crystal violet to quantify biofilm. Data from six biological assays. Statistical analyses using One-Way ANOVA followed by Dunn’s multiple comparisons test. ** p<0.01; *** p<0.001, when not indicated, n.s. (not significant). **E.** Growth of wild-type and mutant strains in LB medium. **F.** Growth of wild-type and mutant strains under iron deficiency by addition of 150 μM DP to the LB medium. The curves were determined by measuring the OD_600nm_ of the cultures during the first eight hours (1-hours intervals) and at 24 hours (items E and F).

### VitR controls many processes by acting as a direct repressor of *vioS*

To identify VitR-regulated genes, we performed RNA-seq on WT and Δ*vitR* strains grown in LB at HCD (Table S2; Figure 4A and B). The gene with the highest expression in Δ*vitR* was *vioS*, which encodes a protein that inhibits violacein production,^33^ possibly by inhibiting the QS regulator CviR through protein-protein interaction.^33^ Many downregulated genes in Δ*vitR* are involved in processes that are regulated by CviR (violacein biosynthesis, proteases, chitinase), suggesting that their altered expression levels in Δ*vitR* is an indirect effect of *vioS* overexpression (Table S2; Figure 4A and B). The *vitR* gene is next to and is divergently transcribed in relation to *vioS* (Figure 1B and 4C). The *vioS* and *vitR* promoter expression was investigated by beta-galactosidase assays (Figure 4C). The expression of *vioS* was fully repressed in the WT strain and entirely de-repressed in the Δ*vitR* mutant in all conditions tested, supporting the RNA-seq data and indicating that VitR represses the *vioS* expression. Iron and Fur have little or no effect on *vioS* expression (Figure 4C). The *vitR* expression levels decreased under iron limitation in the WT, and in Δ*vitR* and Δ*fur,* regardless of iron levels (Figure 4C). These data show that VitR activates itself and is also activated by Fur under iron sufficiency conditions. We investigated whether the purified VitR protein binds to the intergenic region between *vioS* and *vitR* by EMSA assay (Figure 4D). We observed DNA binding starting at 10 nM of VitR, with complete protein binding occurring at 25 nM. This binding was specific, as demonstrated by a competition assay using a nonspecific probe as control (Figure 4D). Altogether, our data indicate that VitR activates its own expression and represses *vioS* by binding directly to their promoters. To confirm that VitR operates via *vioS*, we generated Δ*vioS* and Δ*vitR*/*vioS* mutant strains. While Δ*vitR* had lower violacein production, biofilm formation, and proteolytic activity and an increased siderophore halo, for Δ*vioS* and Δ*vitR*/*vioS* all these phenotypes were similar to those observed in the WT strain (Figure 5). These data demonstrate that the Δ*vitR* phenotypes are exclusively attributed to the de-repression of *vioS* in this mutant.

**Figure 4.**
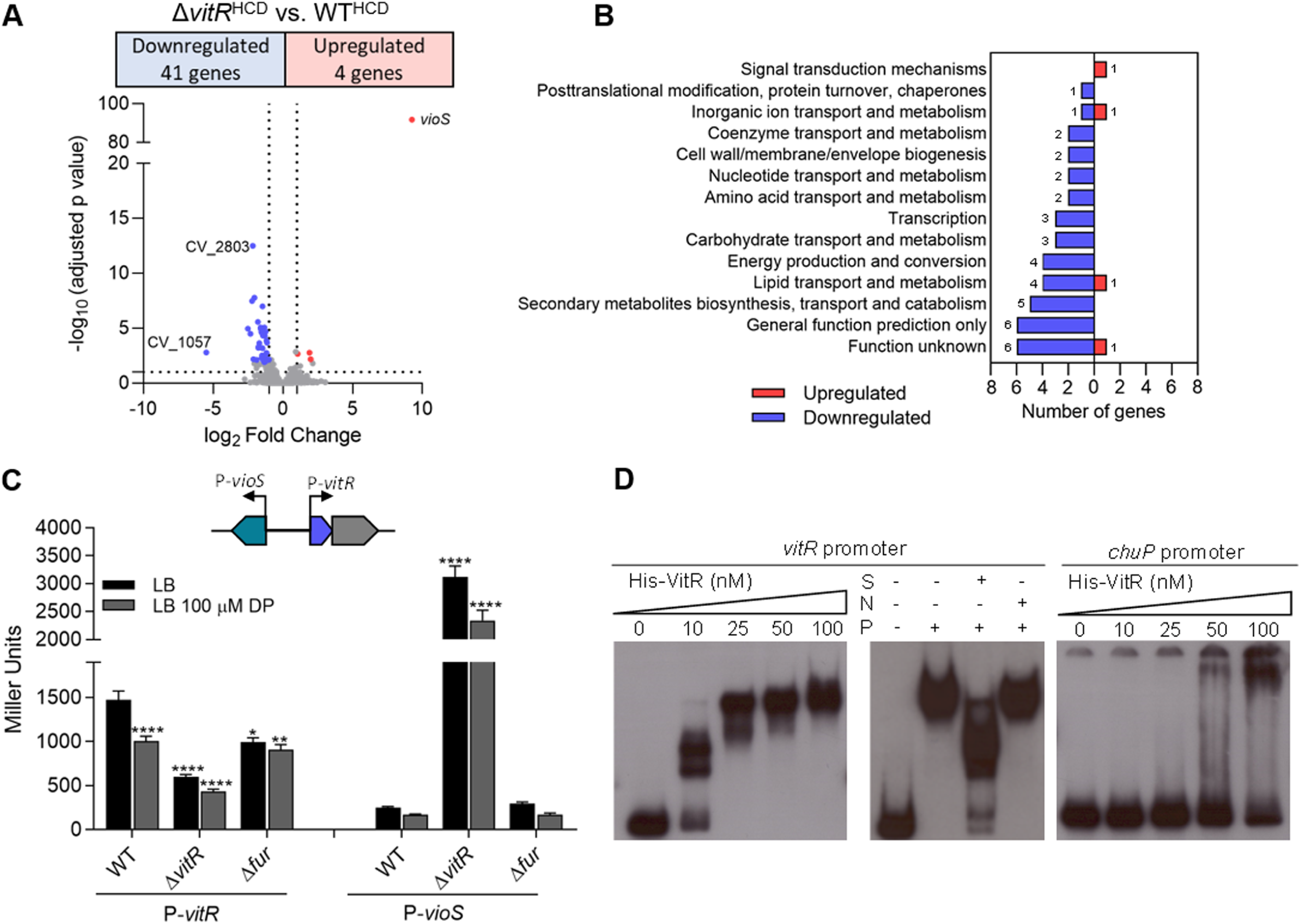
Global expression analysis to identify the VitR regulon in *C. violaceum*. **A.** Volcano plot from RNA-seq data with the distribution of differentially expressed genes in the *vitR* vs. WT comparison. RNA-seq was performed on three biological replicates from bacteria cultured in LB at high cell density (HCD). **B.** Functional categorization of genes regulated by *vitR*. **C.** Expression of the promoter region of the *vitR* and *vioS* genes in the indicated strains. All strains containing the constructs with the indicated promoter were grown until OD_600nm_ 0.6-0.8 and were treated or not for 1 hour with 100 µM of DP. Data from six biological assays. Statistical analyses using Two-way ANOVA followed by Sidak’s multiple comparisons test. ****, p<0.0001, when not indicated, n.s. (not significant). **D.** EMSA with specific (*vioS*/*vitR)* and nonspecific (*chuP*) promoter regions to verify direct binding by the VitR regulator. S-specific unlabeled probe; N-non-specific unlabeled probe; P-50 nM His-VitR protein.

**Figure 5.**
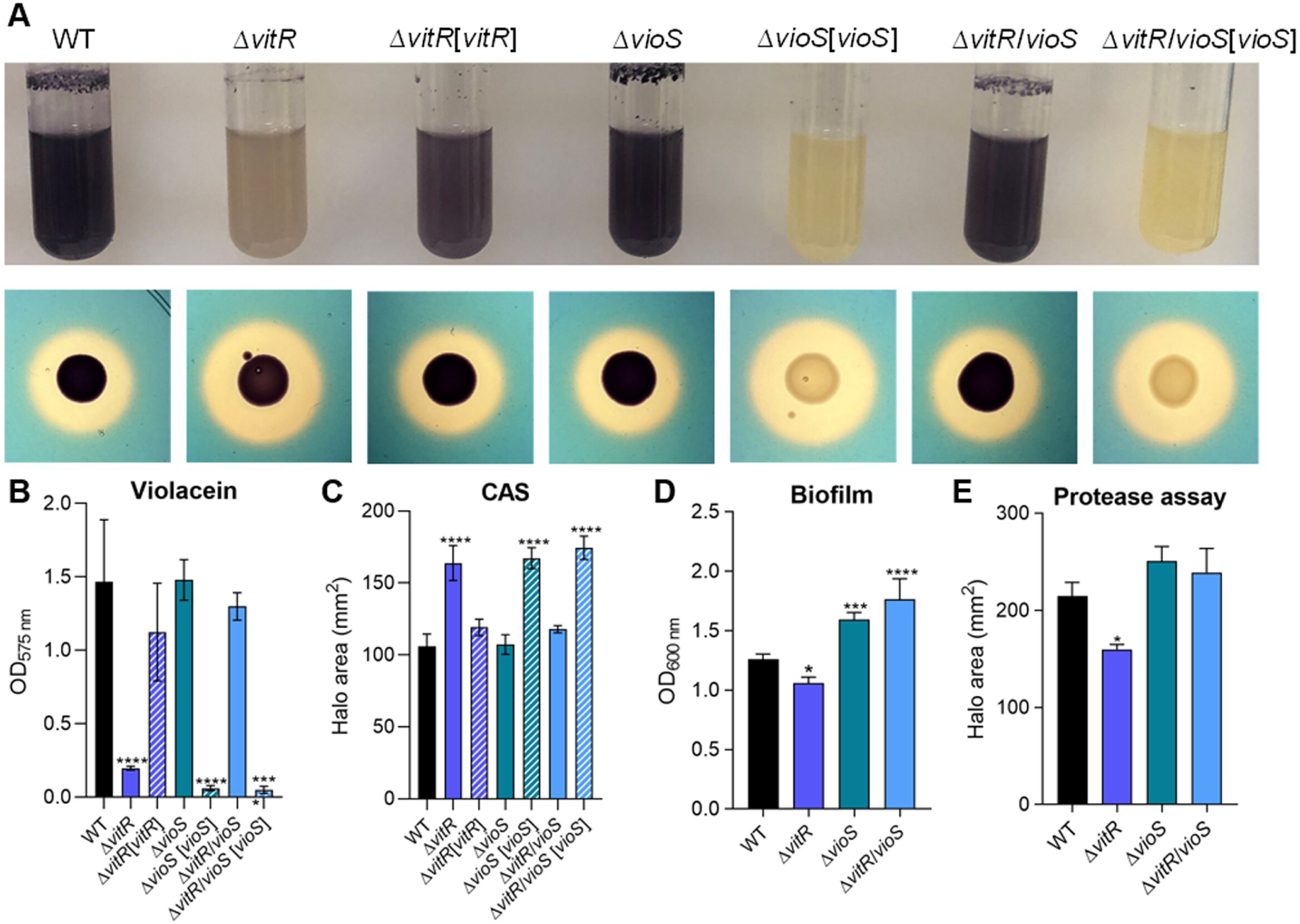
Phenotypic characterization of the Δ*vioS* and Δ*vitR*/*vioS* mutants. **A.** Growth of the indicated strains in LB medium to verify the production of violacein and CAS tests showing the production of siderophores (orange halos) by the indicated strains inoculated on PSA-CAS plates. **B.** Quantification of violacein production of the indicated strains. The strains were grown in LB medium under agitation at 37°C for 24 hours. After incubation, 500 µL of culture were homogenized with 500 µL of acetone. After centrifugation, OD_575nm_ was measured for quantification of violacein. Data from three biological assays. ****p < 0.0001; when not indicated is not significant. One-way ANOVA followed by Tukey’s multiple-comparison test. **C.** Measurement of the CAS halo areas of the indicated strains. Data from three biological assays. Statistical analyses using One-way ANOVA followed by Dunnett’s multiple comparisons test. **** p<0.0001, when not indicated, n.s. (not significant). **D.** Biofilm assay of the indicated strains. The strains were grown in LB medium for 24 hours and the assay was performed with crystal violet to quantify biofilm. Data from six biological assays. Statistical analyses using One-way ANOVA followed by Dunnett’s multiple comparisons test. * p<0.05; *** p<0.001; **** p<0.0001, when not indicated, n.s. (not significant). **E.** Protease tests showing the production and secretion of proteases in M9 plates supplemented with 1.5% of powdered milk. Measurement of the proteases halo areas of the indicated strains. Data from three biological assays. Statistical analyses using One-way ANOVA followed by Dunnett’s multiple comparisons test. * p<0.05; *** p<0.001; **** p<0.0001, when not indicated, n.s. (not significant).

### The CviIR QS system controls siderophore homeostasis in *C. violaceum*

Among the transposon-mutant strains with increased siderophore halos, 51 strains (50%) had insertions in the coding region or in the promoter of the *cviR* gene (CV_4090), which encodes the regulator of the *C. violaceum* CviIR QS system (Table S1; Figure 1D). To confirm that the CviIR QS system controls siderophores, we performed the PSA-CAS assays using *cvi* null mutant strains. As expected, the Δ*cviR* and Δ*cviI* mutants had increased siderophore halos compared to the WT strain, with all the complemented strains having the phenotype reversed (Figure 6A and B). To verify whether the increased siderophore halos in the *ΔcviR* strain were related to a specific siderophore, we generated insertion mutants in each of the NRPS genes (*cbaF* and *vbaF*) using the Δ*cviR* mutant as a background. In both double-mutants, the size of the siderophore halos was slightly smaller compared to that in the Δ*cviR* strain (Figure 6A and B), suggesting that the CviIR QS system of *C. violaceum* affects the homeostasis of both siderophores. Considering that the deletion of either *cviR*, *vitR*, or *airR* led to increased siderophore halos, we tested whether these transcription factors regulate the expression of the *cviR* promoter in an iron-dependent manner. The beta-galactosidase assays revealed that (i) there is no difference in *cviR* expression under iron deficiency; (ii) CviR is not self-regulated; (iii) VitR does not regulate *cviR* expression, which agrees with VitR acting on CviR via VioS; and (iv) under these conditions, AirR does not regulate *cviR* (Figure 6C).

**Figure 6.**
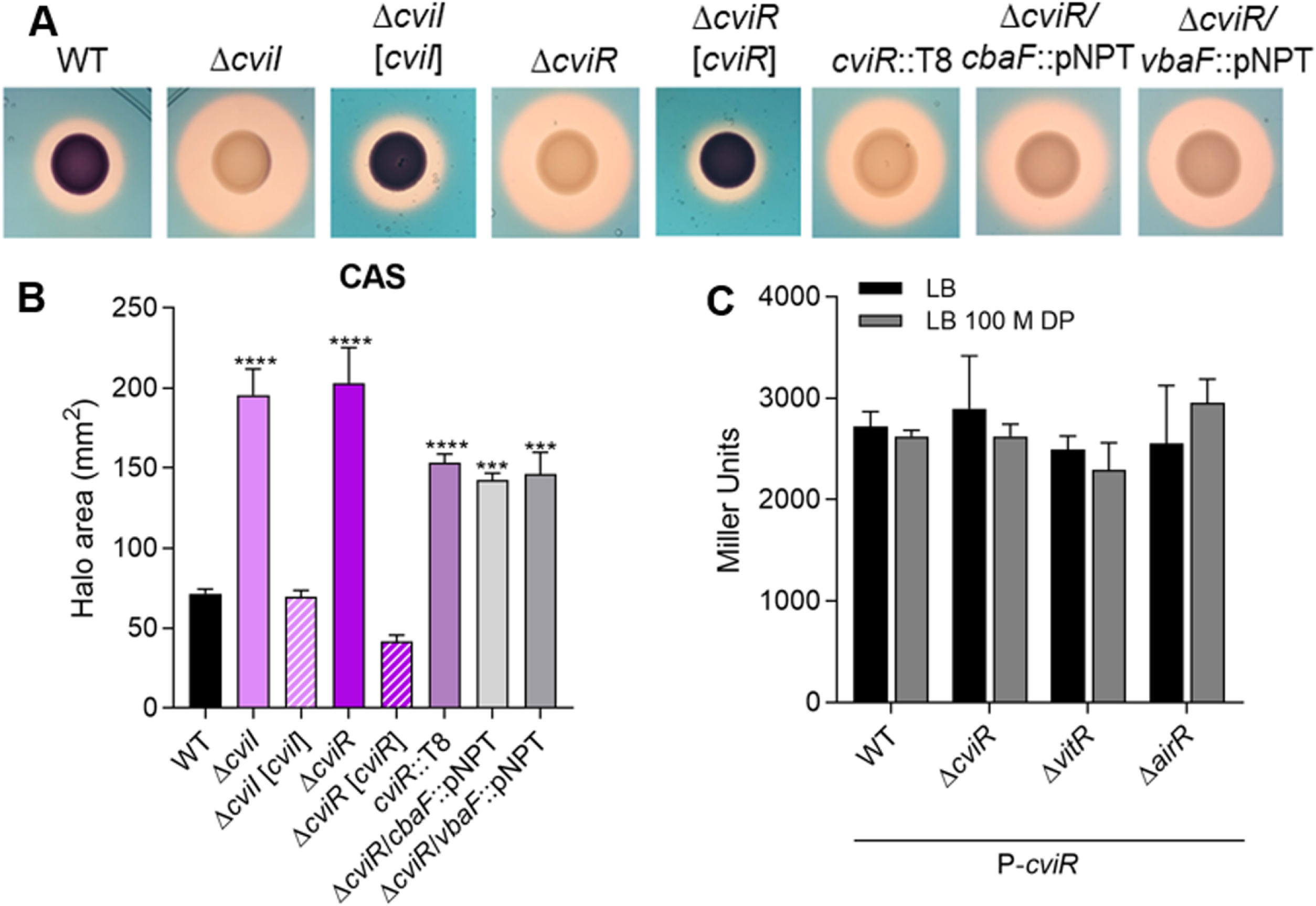
The CviIR QS system regulates siderophore activity in *C. violaceum*. **A.** CAS tests showing the production of siderophores (orange halos) by the indicated strains inoculated on PSA-CAS plates. **B.** Measurement of the CAS halo areas of the indicated strains. Data from three biological assays. Statistical analyses using One-Way ANOVA followed by Dunnett’s multiple comparisons test. *** p<0.001; **** p<0.0001, when not indicated, n.s. (not significant). **C.** Expression of the promoter region of *cviR* in the indicated strains. All strains containing the constructs with the indicated promoter were grown until OD_600nm_ ∼3.0 and were treated or not for 1 hour with 100 µM of DP. Data from three biological assays.

### CviR regulates CviI-dependent and CviI-independent regulons

Given the connections of VitR and AirR with CviR and the shared phenotype of increased siderophore halos in the mutants of these transcription factors, we speculated that the CviIR QS system regulates genes involved in siderophore/iron homeostasis. Despite the many studies on the CviIR QS system,^23,24,25,26,27,34^ the global repertoire of genes regulated by CviI and CviR remains unknown in *C. violaceum*. To compare the transcriptome profiles of the WT, *ΔcviR*, and *ΔcviI* strains, we performed RNA-seq from these strains grown in LB on HCD. There were more differentially expressed genes (DEGs with more than two-fold changes) in the absence of *cviR* (956 DEGs) (Table S2; Figure 7A) than it did in the absence of *cviI* (470 DEGs) (Table S2; Figure 7B). CviR/CviI had a global transcriptional impact, regulating most cell processes (Figure S1A and C). Using RT-qPCR, we validated the expression profile of several up and down-regulated genes in *ΔcviR* and *ΔcviI* (Figure 7C and D, Figure S1B and D). Most of the genes regulated by *cviI* were also regulated by *cviR* (84%), while fewer genes regulated by *cviR* depended on *cviI* (41%) (Figure 7E and F), suggesting that CviR regulates many genes without its CviI-produced canonical autoinducer. We observed that almost all VitR-regulated genes belong to the CviIR regulons (80%), with *vioS* being the only gene that was exclusively repressed by VitR (Figure 7E and F). These data support the hypothesis that VitR acts upstream to the CviIR system via the VioS protein. To evaluate if VioS inhibits CviR by protein-protein interaction, we performed a double-hybrid yeast assay. Our data indicates that VioS directly interacts with the CviR protein (Figure S2).

**Figure 7.**
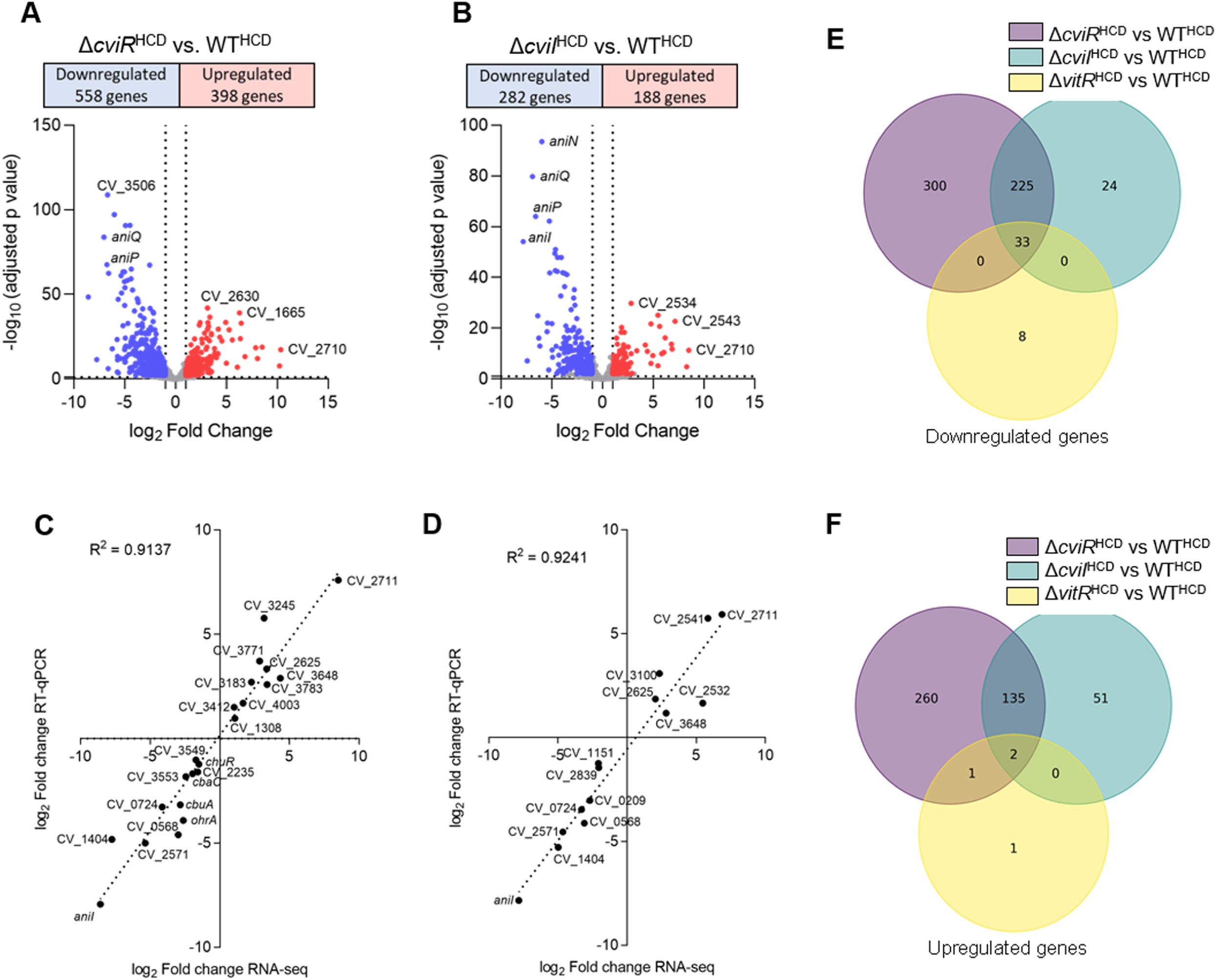
Genome-wide analysis of CviI and CviR-regulated genes in *C. violaceum*. **A.** Volcano plot from RNA-seq data with the distribution of differentially expressed genes in the Δ*cviR* vs. WT comparison. RNA-seq was performed in three replicates from bacteria cultured in LB at high cell density (HCD). **B.** Volcano plot from RNA-seq data with the distribution of differentially expressed genes in the Δ*cviI* vs. WT comparison. RNA-seq was performed in three replicates from bacteria cultured in LB at high cell density (HCD). **C.** Correlation of differentially expressed genes in Δ*cviR*. The log_2_ fold changes obtained from the RNA-seq data were plotted against the log_2_ fold changes determined by RT-qPCR for the indicated genes. **D.** Correlation of differentially expressed genes in Δ*cviI*. The log_2_ fold changes obtained from the RNA-seq data were plotted against the log_2_ fold changes determined by RT-qPCR for the indicated genes. **E-F.** Venn diagrams showing the overlap and unique subset of genes whose expression was lower (E) or higher (F) in the Δ*vitR*, Δ*cviI* and Δ*cviR* strains at high cell density (OD 4.0). Purple circles represent differentially expressed genes in the Δ*cviR* strain compared to the WT^HCD^ strain. Blue circles indicate differentially expressed genes in the Δ*cviI* strain compared to the WT^HCD^ strain. Yellow circles indicate differentially expressed genes in the Δ*vitR* strain compared to the WT^HCD^ strain. RNA-seq was performed in three replicates per strain and per condition.

### Expression of CviR and CviI-regulated genes according to cell density

To reveal whether the expression of CviR and CviI-regulated genes change according to cell density, we compared the RNA-seq data from the WT at HCD (this work) with that from the WT at LCD grown under the same conditions (unpublished data) (Table S2). As expected, most CviR and CviI-regulated genes (68%) had their expression levels altered according to cell density (Figure 8A).

**Figure 8.**
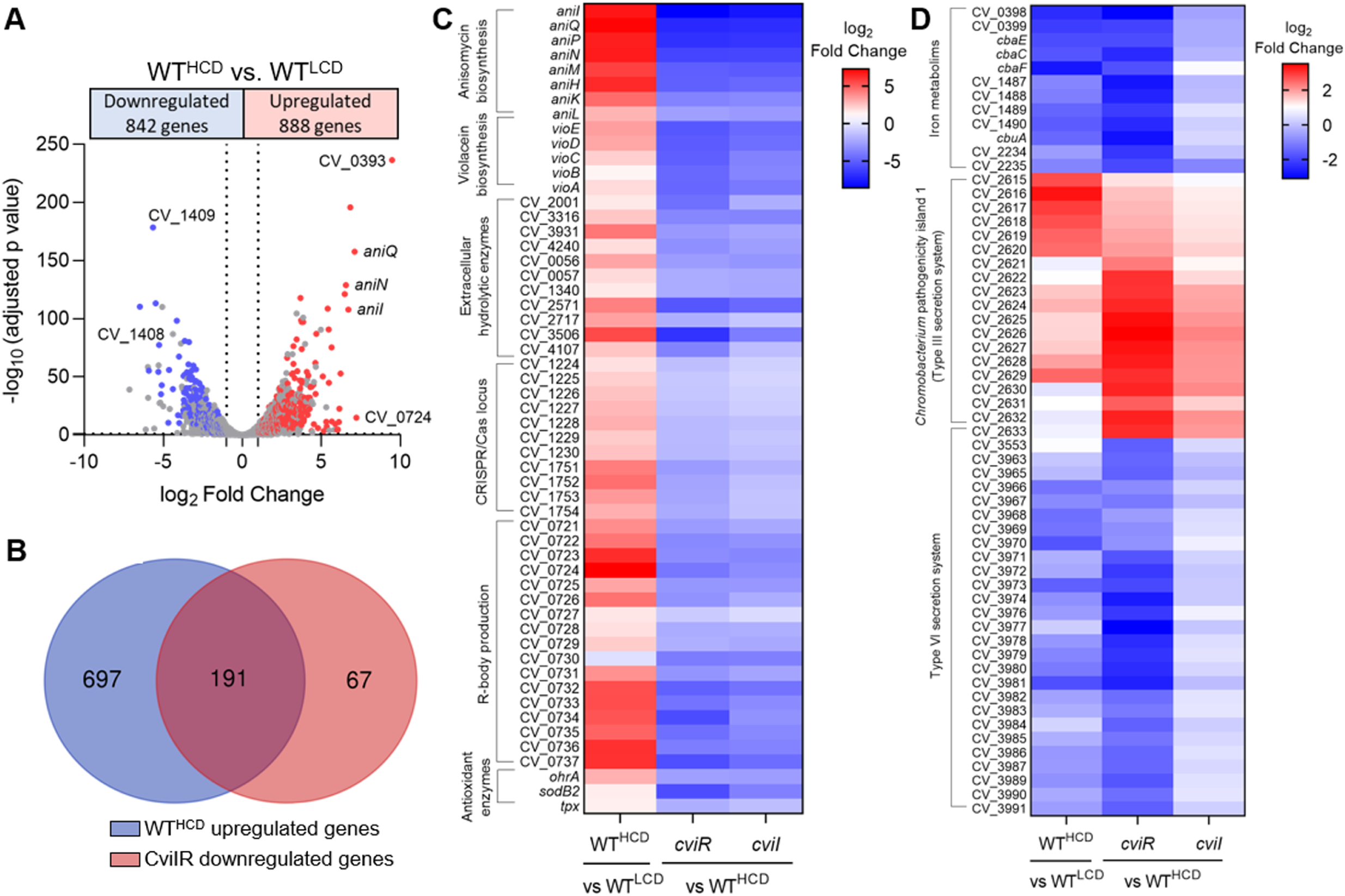
*C. violaceum* has distinct CviR and CviI regulons. **A.** Volcano plot with the distribution of differentially expressed genes in the WT^HCD^ vs WT^LCD^ comparison. Colored dots highlight the genes that were also differentially expressed in the mutants of the CviIR QS system. *C. violaceum* ATCC 12472 was grown in LB medium to OD ∼1.0 for a low cell density (LCD) condition and to OD ∼4.0 for a high cell density (HCD) condition. RNA-seq was performed in three replicates. **B**. Venn diagrams showing the overlap and unique subset of genes upregulated in HCD and downregulated in the mutants of the CviIR QS system. The blue circle represents upregulated genes in the WT^HCD^ strain. The red circle indicates downregulated genes in the Δ*cviR* and Δ*cviI* strains. **C.** Heatmap showing a subset of the genes activated by the CviIR QS system at HCD. **D.** Heatmap showing a subset of the genes regulated by the CviR, regardless of cell density and Cvil. Comparison of WT^HCD^ vs WT^LCD^, Δ*cviR* vs WT^HCD^, and Δ*cviI* vs WT^HCD^.

### Genes related to classical QS-dependent processes are activated by CviIR at high cell density

To identify the processes activated by CviR/CviI at HCD, we focused on 258 genes downregulated in both *ΔcviR* and *ΔcviI* (Figure 7E). Of these, 191 genes were also upregulated in WT at HCD (Figure 8B). This group of 191 genes includes genes encoding a lectin (CV_1744), many extracellular hydrolytic enzymes (1 collagenase, 3 chitinases, 7 protases), and clusters of antibiotic biosynthesis (*vioABCDE* for violacein and *aniIQPMNHKL* for anisomycin) (Figure 8C) that are known QS-associated processes in *C. violaceum*.^25,26^ Also included in this group, there are several large gene clusters (CV_1395 to CV_1407, CV_1541 to CV1547, CV_2798 to CV_2804, CV2831 to CV_2837, and CV_3940 to CV_3961), which may be related to the production of new small bioactive metabolites that were previously detected but not identified in a metabolome analysis of *C. violaceum*.^26^ Remarkably, two CRISPR/Cas loci (CV_1224 to CV_1230 and CV_1751 to CV_1754), the gene clusters for CvP4 phage (some genes of CV_2114 to CV_2150) and R-body production (CV_0721 to CV_0737), and genes encoding antioxidant enzymes (*ohrA*, *sodB2*, *tpx*) were also activated by CviIR at HCD (Figure 8C).

### Genes related to iron/siderophore uptake are activated by CviR at low cell density

Our RNA-seq data revealed that many genes related to iron/siderophore acquisition were downregulated in *ΔcviR* but their expression levels were almost unaffected in *ΔcviI*. Consistent with a mechanism of CviR activation at LCD, most of these genes were expressed at higher levels at LCD than at HCD (Figure 8D). Such genes encode transporters for iron acquisition (*feoB*, *exbBD*), including iron bound to the siderophores chromobactin (*cbuA* and CV_1487-88-89), viobactin (CV_2234-35) and heme (*chuR*). Therefore, the increased siderophore halos in *ΔcviR* seems to be related to an impaired siderophore uptake. These data suggest that in addition to being regulated by Fur in response to iron levels, as reported by Santos et al.,^30^ the genes involved in siderophore-mediated iron acquisition are also activated by CviR at LCD, which must optimize iron uptake. Many other processes beyond the scope of this study were regulated by the CviIR QS system. For instance, almost every gene of a large cluster encoding the type VI secretion system (T6SS) was downregulated in *ΔcviR*, and genes encoding the Cpi1 type III secretion system (T3SS) were upregulated in *ΔcviR* and *ΔcviI* (Figure 8D).

## Discussion

Siderophores enable bacteria to survive in iron scarcity, an environmental condition commonly encountered by pathogenic and free-living bacteria. However, siderophore synthesis and utilization must be finely regulated to avoid superfluous production, toxicity, and unwanted use of siderophores by non-producing organisms.^16,35,36^ In this study, we used an unbiased transposon mutagenesis approach to identify novel regulatory systems involved in siderophore-mediated iron homeostasis in *C. violaceum* (Table S1), a bacterial pathogen that relies on endogenous siderophores to infect mammalian hosts.^28^ Among the identified regulatory systems, we characterized a regulatory cascade involving the transcription factor VitR, the two-component system AirSR, and the QS system CviIR (Figures 1 and 9). This cascade allows *C. violaceum* to tailor the expression of siderophore-mediated iron acquisition genes according to cell density, adding a novel layer of regulation to the already known iron level-based Fur-mediated repression.^30^

**Figure 9.**
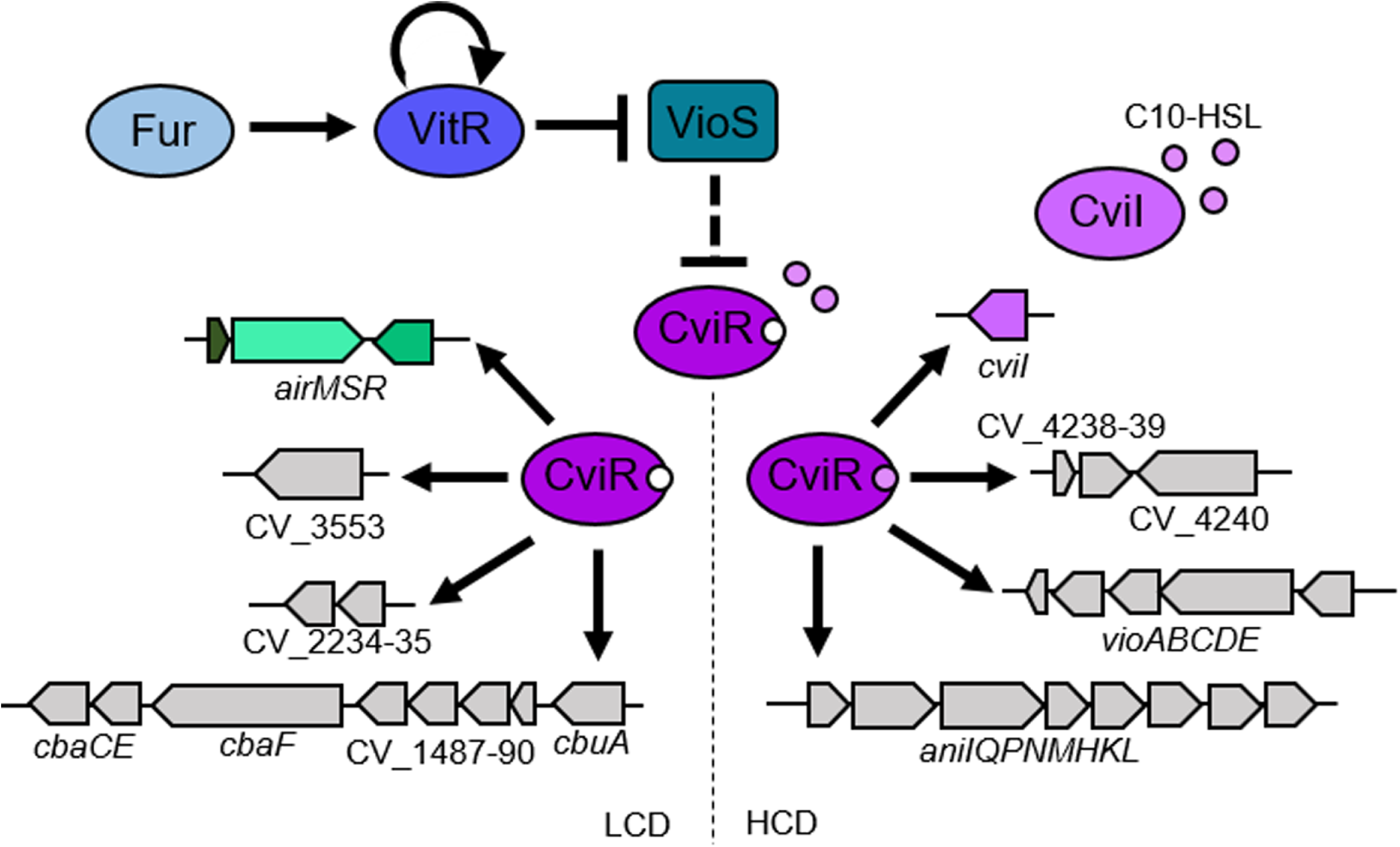
A regulatory cascade involving the CviIR QS system regulates siderophores in *C. violaceum*. The VitR regulator under iron sufficiency is self-activated and activated by Fur. VitR is a dedicated repressor of *vioS*, which encodes a protein that inhibits CviR through protein-protein interaction (dashed line). At high cell density in the presence of the CviI autoinducers, the CviR regulator activates the production of AHLs (*cviI*), violacein (operon *vioABCDE*), anisomycin (operon *aniQPNMHKL*), and proteases (CV_4240). In a CviI-independent mechanism, the CviR regulator activates the AirMSR system and the genes necessary for the uptake of siderophores (CV_3553, CV_2234, CV_2235, *cbaF*, *cbaCE*, CV_1487-40 e *cbuA*) at low cell density in *C. violaceum*.

In this study, we characterized VitR as a novel transcription factor that controls siderophore, violacein, and biofilm formation in *C. violaceum* (Figure 3). Our data provide evidence that VitR operates upstream of the CviIR QS system by acting as a direct repressor of *vioS*, as follows: (i) VitR binds to the intergenic region where divergent promoters of the *vioS* and *vitR* genes are found, repressing *vioS* and activating its own expression (Figure 4); (ii) all phenotypes of Δ*vitR* mutant were rescued in a double Δ*vitR*/*vioS* mutant (Figure 5); (iii) almost all VitR-regulated genes belong to the CviIR regulons (Figures 4 and 7E and F). In addition, we demonstrated a direct interaction between VioS and the QS regulator CviR (Figure S2), corroborating a previous hypothesis that VioS inhibits CviR through protein-protein interaction.^32,33^ Therefore, unlike other iron-sensing transcription factors that in response to iron directly regulate large regulons, such as XibR and VgrR in *X. campestres*,^12,37^ VitR exerts its effects as a dedicated local repressor of *vioS*, and its connection with iron seems to be indirect via Fur-mediated *vitR* regulation. The signal that releases VitR from DNA to trigger *vioS* expression remains to be determined. VitR belongs to the Cro superfamily, XRE family of transcription factors. DNA binding of XRE family members can be antagonized by small molecules (BzdR regulator) or by DNA mimic proteins (NHTF regulator).^38,39,40^ Thus, we hypothesize that CV_1058, a protein of unknown function, could inhibit the DNA-binding activity of VitR through a protein-protein interaction mechanism.

Our data indicate that CV_0535-36-37 plays a role in the regulation of siderophores in *C. violaceum* ATCC 12472 (Figures 1 and 2). This system has been characterized in *C. violaceum* ATCC 31532 as an antibiotic-induced system (*airMSR*) acting through the CviIR signaling pathway to activate violacein production during *C. violaceum* competition with *Streptomyces* spp.^32^ We tested violacein production in *airMSR* mutants in *C. violaceum* ATCC 12472, but the phenotypes that have been described in *C. violaceum* ATCC 31532 were not observed (data not shown). Thus, the CV_0535-36-37 system in *C. violaceum* ATCC 12472 appears to sense signals other than antibiotics.

An important finding of our study was that the CviIR QS system is involved in the regulation of siderophores in *C. violaceum*, since the mutation of the *cviIR* genes led to increased siderophore halos on PSA-CAS plates (Figure 6). In agreement with our data, it has been described that mutation of QS systems leads to increased siderophores in *Burkholderia ambifaria*, *Pseudomonas chlororaphis*, and *Vibrio vulnificus*.^14,15,17^ To understand how the CviIR QS system controls siderophore levels in *C. violaceum*, we identified the entire repertoire of CviR and CviI-regulated genes (Figure 7) and checked whether their regulation was cell-density dependent (Figure 8). Surprisingly, we found that CviR regulates CviI-dependent and CviI-independent regulons, suggesting that CviR can act regardless of its canonical CviI autoinducers, which in *C. violaceum* ATCC 12472 are several long-chain AHLs.^23,26^ For instance, almost all genes of a large cluster encoding the T6SS were downregulated in *ΔcviR* but not in *ΔcviI,* which is consistent with our previous data that CviR, but not CviI, is required for *C. violaceum* T6SS-mediated competition.^41^ Dissimilar phenotypes and regulons were also described in the *Pseudomonas aeruginosa* RhlI/R QS system,^42^ and in this case an alternative PqsE-produced ligand accounts for the expression of RhlR dependent genes in the absence of RhlI.^43^ As expected, genes encoding classical QS-dependent processes described in *Chromobacterium spp.* were activated at high cell density by both CviI and CviR (Figure 8), including those for extracellular enzymes like proteases and chitinases, and for biosynthesis of the antibiotics violacein and anisomycin.^24,25,26,44^

Siderophores are considered public goods, like many extracellular enzymes.^45,46^ However, the pattern of QS-mediated regulation of genes related to iron/siderophore acquisition was distinct than what was observed for extracellular enzymes, since siderophore genes were more expressed at LCD than at HCD and were downregulated in *ΔcviR* but not in *ΔcviI* (Figure 8). These results suggest that *C. violaceum* boosts its capacity to acquire iron via siderophores at LCD via CviR activation. A similar QS regulatory strategy has been described for the global QS regulator LuxT, which regulates siderophores in *Vibrio harveyi*.^47^ In this bacterium, the same QS-regulated siderophore cluster produces cell-associated and soluble siderophores to optimize iron uptake according to the bacteria life stages.^16^ It has been described in *Paracoccus denitrificans* that a QS system leads to a shift from TonB-dependent to TonB-independent iron uptake strategies during biofilm formation.^48^

Future studies should investigate the role of the CviIR QS system in the pathogenesis of *C. violaceum* infecting mammals, as this system has been investigated in invertebrate models.^26,34^ Also, a detailed analysis of the chemical structure of siderophores chromobactin and viobactin may provide insight into their role in different *C. violaceum* life stages.

## Materials and Methods

### Bacterial strains, plasmids, and growth conditions

All the strains and plasmids used in this work are described in Table S3. *E. coli* strains were cultured in Luria-Bertani (LB) medium and *C. violaceum* strains were cultured in LB medium or M9 minimal medium supplemented with 0.1% casein hydrolysate. When necessary, cultures were supplemented with kanamycin (50 μg/mL), ampicillin (100 μg/mL), nalidixic acid (4 μg/mL), gentamicin (40 μg/mL for *C. violaceum* or 20 μg/mL for *E. coli*) or tetracycline (5 μg/mL in liquid medium for *C. violaceum*, 10 ug/mL in agar plates for *C. violaceum* or 12 μg/mL for *E. coli*). Iron deficient conditions were obtained by supplementation with 2,2^’^-dipyridyl (DP) (Sigma) as previously defined for *C. violaceum*.^28^ *Saccharomyces cerevisiae* AH109 strain was cultured in yeast peptone dextrose adenine (YPDA) medium.^49^

### Generation and screening of a transposon mutant library

To obtain transposon mutants in *C. violaceum*, the IS*lacZ*/hah (T8) transposon present in the pIT2 plasmid was used.^31^ We have validated this transposon to generate mutants in *C. violaceum* using as background a spontaneous nalidixic acid resistant mutant (CV^NALR^).^30^ To obtain several random insertion mutants, the *C. violaceum* CV^NALR^ strain was conjugated with *E. coli* SM10λpir carrying the pIT2 vector. A library with approximately 10,000 mutants was organized in 96-well plates and frozen at -80 °C. These *C. violaceum* mutants were screened for altered halos in the siderophore-indicative PSA-CAS plates. Mutants with decreased or increased halos were selected and the transposon insertion site was identified by semi-degenerate PCR (Table S4) followed by Sanger sequencing as described.^30,31^

### Construction of *C. violaceum* mutant strains

Null-mutant strains were derived from the wild-type strain ATCC 12472 and generated by allelic exchange (in-frame null deletion) as previously described.^28,50^ Primers used for cloning, sequencing and mutant confirmation are listed in Table S4.

### Construction of complemented strains

Null mutants were trans-complemented with the wild-type copies of the genes containing their promoter regions cloned into the replicative plasmids pMR20 or pSEVA. Primers used for cloning are listed in Table S4.

### Siderophore production assay

Measurement of siderophore production was performed by the universal Chrome Azurol S (CAS) agar plate assay^51^ with replacement of the MM9 medium by peptone-sucrose agar (PSA).^28,52^ Ten microliters of *C. violaceum* cultures were spotted on PSA-CAS agar plates, and siderophore production was evaluated by orange halos that appeared after incubation for 24 hours at 37°C. The area of the halos was measured using the ImageJ program. All experiments were performed in three biological replicates.

### Static biofilm

For the biofilm assay, the strains were grown in LB medium from an OD_600nm_ of 0.01 in glass tubes and incubated at 37 °C without shaking for 24 hours. After incubation, the cultures were washed, and the biofilm was stained with 0.1% crystal violet. After washes, the biofilm was resuspended in 1 mL of 100% ethanol and the OD_600nm_ was measured. Experiments were performed in six replicates.

### Growth curves

To evaluate the growth of the mutant strains over time, growth curves were performed. For this experiment, the wild-type and mutant strains were initially cultivated in LB medium overnight and the cultures were diluted to an OD_600nm_ of 0.01 in 4 mL of LB medium and incubated under agitation (250 rpm). Growth was determined by measurement of the OD_600nm_ for the 8 initial points, in addition to the 24 hours point. For an iron deficient condition, LB medium was supplemented with 150 µM of DP. The experiments were performed in three biological replicates.

### Violacein production

To analyze the violacein production of the different strains, an initial cultivation was performed in LB medium at 37 °C overnight, and then cultures were diluted to an OD_600nm_ of 0.01 and incubated at 37 °C under agitation. The cultures were photographed after 24 hours to verify violacein production. To quantify the violacein production, 500 µL of the cultures were mixed with 500 µL of 100% acetone. Tubes were vortexed for 30 seconds and centrifuged for 5 minutes at 13000 rpm. The organic phase, containing violacein, was quantified in a spectrophotometer at a wavelength of 575 nm. The experiments were performed in three biological replicates.

### Protease assay

To verify the presence of proteases, the wild-type and mutant strains were grown overnight in M9 medium at 37 °C. Ten microliters of these cultures were plated on the surface of M9 medium supplemented with 1.5% powdered milk replacing the casein hydrolysate. The plates were incubated at 37°C for 24 hours and the halos produced were measured using the ImageJ program. The experiments were performed in three replicates.

### RNA purification and RNA-seq

Total RNA of the wild-type, Δ*vitR*, Δ*cviI*, and Δ*cviR* strains were extracted from three independent biological replicates. The bacterial strains were grown in LB medium at high cell density (OD_600nm_ ∼ 4,0), and the RNA samples were extracted using TRIzol reagent and purified using Direct-zol RNA Purification Kit (Zymo Research), following manufacturer’s instructions. Purified RNAs were sent to NGS Soluções Genômicas for RNA sequencing (RNA-seq) (https://ngsgenomica.com.br). The RNA integrity was verified using a 2100 Bioanalyzer instrument (Agilent Technologies). Depletion of rRNA and cDNA library preparation were performed using lllumina Stranded Total RNA Prep with Ribo-Zero Plus (Illumina). The cDNA libraries were quantified by qPCR followed by sequencing in a NextSeq2000 equipment (Illumina).

### Bioinformatic analysis of RNA-seq data

The raw data were processed using the frtc pipeline available at [https://github.com/alanlorenzetti/frtc/].^53^ Briefly, the quality of the reads was checked using Rqc,^54^ the adapters were trimmed, and the remaining low-quality ends (Q<30) were removed using Trimmomatic.^55^ The trimmed reads were aligned against the reference genome (*C. violaceum* ATCC 12472, genome assembly ASM770v1) using HISAT2.^56^ Differential expression analysis was performed using the scripts available at https://github.com/alanlorenzetti/ccrescentus_RNASeq_analysis.^57^ The read counts were performed using GenomicAlignmets^58^ and differential expression analysis using DESeq2^59^ with the following cluster design: Δ_*vitR*_HDC vs. WT_HDC, Δ_*cviI*_HDC vs. WT_HDC, Δ_*cviR*_HDC vs. WT_HDC, and WT_HDC vs. WT_LDC). Genes with log2 fold change ≥ 1 or ≤ -1 and adjusted p-value < 0.01 were considered differentially expressed. Functional categorization was performed using the Clusters of Orthologous Groups (COG) with some manually added annotations based on previous lab work. The raw data for the manuscript: RNA sequencing results of *C. violaceum* wild-type (WT), Δ*vitR*, Δ*cviI*, and Δ*cviR* strains at high cell density (accession number PRJNA1003908) and RNA sequencing results of *C. violaceum* wild-type (WT) at low cell density (accession number PRJNA1006746) are available on Sequence Read Archive (SRA) or upon request from the lead contact Prof. José Freire da Silva Neto.

### Construction of transcriptional *lacZ* fusions and β-galactosidase assay

The region upstream of the genes of interest was amplified by PCR with proper primers (Table S4) and cloned into the pGEM-T easy plasmid (Promega). The insert was removed by digestion and subcloned into the pRK*lacZ*290 vector to generate transcriptional fusions to the *lacZ* gene. *C. violaceum* cultures harboring these reporter plasmids were grown in different conditions: (i) LB medium until OD_600nm_ 0,8-1,0; (ii) LB medium until OD_600nm_ 0,6-0,8 and treated or not with 100 µM of DP for one hour. Next, cells were assayed for β-galactosidase activity based on a previously described protocol.^30^

### Gene Expression by RT-qPCR

The *C. violaceum* wild-type, Δ*cviR*, and Δ*cviI* strains were grown in LB medium until high cell density (OD_600nm_ ∼4.0). Total RNA was extracted and purified as described above. Two micrograms of total RNA from each sample were converted to cDNA using the High-Capacity cDNA Reverse Transcription kit (Thermo Fisher Scientific). Quantitative PCR (qPCR) reactions were performed using the PowerUp™ SYBR™ Green Master Mix (Thermo Fisher Scientific), the specific primers (Table S4), and 0.5 μL of cDNA. The relative expression was calculated by the 2^-ΔΔCt^ method.^60^ Data from three biological replicates were normalized by an endogenous control (*minD* gene) and a reference condition (WT).

### Expression and purification of VitR

The coding region of the *vitR* gene was PCR-amplified (Table S4) and cloned into the pET15b vector (Table S3). The recombinant histidine-tagged protein was overexpressed in *E. coli* BL21(DE3) by induction with 1 mM isopropyl-D-thiogalactopyranoside (IPTG) for 2 hours at 37°C in LB medium. After induction, the soluble fraction containing the His-VitR protein was purified using NTA-resin affinity chromatography in phosphate buffer according to manufacturer’s recommendations (Qiagen). After concentration (Vivaspin 6 Concentrator, Sartorius Stedim Biotech) and desalting (PD 10 Desalting Columns, GE Healthcare), the purified VitR protein was resolved by 15% SDS-PAGE.

### Electrophoretic mobility shift assay (EMSA)

The promoter regions of the *vitR* and *chuP* genes were amplified by PCR using the oligonucleotides listed in Table S4. These DNA fragments were labeled with [γ-32P]ATP (PerkinElmer) by using T4 polynucleotide kinase (Thermo Scientific) and purified with the NucleoSpin Gel and PCR Cleanup kit (Macherey-Nagel). The DNA binding reactions were performed in interaction buffer (10 mM Tris-HCl [pH 7.5], 40 mM KCl, 1 mM MgCl_2_, 0.1 mg/mL bovine serum albumin, 1 mM DTT, 5% glycerol), 0.1 mg/mL competitor salmon sperm DNA, DNA probes and different concentrations of His-VitR at a final volume of 20 µL. All interaction reactions were incubated at 25°C for 25 min. Next, 3 µL of 50% glycerol were added, and the samples were separated by native 5% polyacrylamide gel electrophoresis in Tris-borate (TB) buffer. Competition assays were performed using 50 nM of His-VitR as described above in the presence of a 10-fold excess of unlabeled specific (promoter region *vitR*) or non-specific (promoter region *chuP*) probes. The gels were dried, and the signal was detected by autoradiography.

### Yeast double-hybrid assay

To verify protein-protein interaction, we performed a double-hybrid assay in *S. cerevisiae* according to Lin and Lai.^49^ Briefly, the coding region of the *vioS* and *cviR* genes were PCR-amplified (Table S4) and cloned into vectors pGADTK7 (prey, fusion with activation domain) and pGBKT7 (bait, fusion with DNA-binding domain), respectively. The constructs were transformed into *S. cerevisiae* strain AH109 and positive colonies were selected on synthetic minimal medium without leucine and tryptophan supplementation (SD-WL). To verify the protein-protein interaction, different clones were grown in minimal synthetic medium without leucine, tryptophan, histidine and adenine (SD-WLHA) supplementation.

### Statistical analysis

Statistical analysis was performed in GraphPad Prism version 8. For the column graphs, the normality test was performed using Shapiro-Wilk’s test and group comparison was performed by one-way Analysis of Variance (ANOVA) followed by multiple comparisons test. For the grouped graphs used in the β-galactosidase assay statistical analysis was performed by Two-way ANOVA followed by multiple comparisons test. Statistically significant p values or other tests that were performed are indicated in the figure’s subtitles.

## Acknowledgments

This research was supported by grants from the São Paulo Research Foundation (FAPESP; grants 2018/01388-6 and 2021/06894-0) and Fundação de Apoio ao Ensino, Pesquisa e Assistência do Hospital das Clínicas da FMRP-USP (FAEPA). During the course of this work, B.B.B., V.M.L., and B.A.P. were supported by fellowships from FAPESP (grants 2018/19058-2; 2020/15268-2; 2022/00308-4, respectively) and CAPES (Coordenação de Aperfeiçoamento de Pessoal de Nível Superior). J.F.S.N. is Research Fellow from CNPq (Conselho Nacional de Desenvolvimento Científico e Tecnológico).

## Author contributions

B.B.B. and J.F.S.N. conceived and designed the experiments. B.B.B. and V.M.L. performed the experiments. B.A.P. and T.K. performed the bioinformatics analysis. B.B.B., V.M.L., and J.F.S.N. wrote the paper and edited the manuscript.

## Supplemental Excel table titles and legends

**Supplemental Table 1. Identification of insertion sites of transposon mutant strains screened for altered siderophore levels.**

**Supplemental Table 2. Genes differentially expressed in RNA-seq analyses. Supplemental Table 3. Bacterial strains and plasmids.**

**Supplemental Table 4. Primers used in this work.**

**Supplemental Figure 1. Validation of differentially expressed genes in Δ*cviI* and Δ*cviR* by RT-qPCR. A.** Functional categorization of genes with altered expression in Δ*cviI*. Abbreviations: NA, no annotation. IT, intracellular trafficking. PT, posttranslational. SM, secondary metabolites. **B.** Validation of differentially expressed genes in Δ*cviI.* **C.** Functional categorization of genes present in the CviR regulon. Abbreviations: NA, no annotation. IT, intracellular trafficking. PT, posttranslational. SM, secondary metabolites. **D.** Validation of differentially expressed genes in Δ*cviR*. **B-D.** cDNA was reverse transcribed from total RNA harvested from the WT, Δ*cviI*, and Δ*cviR* strains grown at high cell density (OD_600_ ∼ 4.0). Expression of the indicated genes are shown as the fold change relative to WT normalized by the endogenous control *minD*. ****, p < 0.0001; when not indicated, not statistically significant. Two-way ANOVA followed by Tukey’s multiple comparison test.

**Supplemental Figure 2. VioS interacts directly with the CviR protein. A.** Scheme showing the mechanism of the Two-Hybrid System (Y2H) in *Saccharomyces cerevisiae*. Two proteins of interest (CviR and VioS) are fused with two different domains of Gal4. The bait protein is fused to the DNA-binding domain of Gal4 (VioS) and the tethered protein is fused to the transcriptional activation domain of Gal4 (CviR). In yeast AH109, transcriptional activation of the reporters (ADE2, HIS3 and MEL1) only occurs in the cell if the bait interacts with the prey, leading to the activation of the GAL UAS promoter by the Gal4 transcription factor. **B.** Two-hybrid assay indicating CviR-VioS interaction. The coding regions of the *vioS* and *cviR* genes were cloned to generate fusion with the activating domain and the DNA-binding domain of Gal4, respectively. The vectors with cloning or empty were inserted into the S. cerevisiae strain AH109 through transformation. Colonies selected in SD-WL medium (Synthetic Dextrose Minimal medium, without addition of tryptophan and leucine) were then tested in SD-WLHA medium (Synthetic Dextrose Minimal medium, without addition of tryptophan, leucine, adenine and histidine) to verify the interaction between bait and prey. The empty vector was used as a negative control.

## References

1. Andrews, S.C., Robinson, A.K., and Rodríguez-Quiñones, F. (2003). Bacterial iron homeostasis. FEMS Microbiol Rev 27, 215–237. 10.1016/s0168-6445(03)00055-x.

2. Krewulak, K.D., and Vogel, H.J. (2008). Structural biology of bacterial iron uptake. Biochim Biophys Acta 1778, 1781–1804. 10.1016/j.bbamem.2007.07.026.

3. Wandersman, C., and Delepelaire, P. (2004). Bacterial iron sources: from siderophores to hemophores. Annu Rev Microbiol 58, 611–647. 10.1146/annurev.micro.58.030603.123811.

4. Faraldo-Gómez, J.D., and Sansom, M.S. (2003). Acquisition of siderophores in Gram-negative bacteria. Nat Rev Mol Cell Biol 4, 105–116. 10.1038/nrm1015.

5. Miethke, M., and Marahiel, M.A. (2007). Siderophore-based iron acquisition and pathogen control. Microbiol Mol Biol Rev 71, 413–451. 10.1128/mmbr.00012-07.

6. Bilitewski, U., Blodgett, J.A.V., Duhme-Klair, A.K., Dallavalle, S., Laschat, S., Routledge, A., and Schobert, R. (2017). Chemical and biological aspects of nutritional immunity-perspectives for new anti-infectives that target iron uptake systems. Angew Chem Int Ed Engl 56, 14360–14382. 10.1002/anie.201701586.

7. Braun, V., and Hantke, K. (2011). Recent insights into iron import by bacteria. Curr Opin Chem Biol 15, 328–334. 10.1016/j.cbpa.2011.01.005.

8. Lee, J.W., and Helmann, J.D. (2007). Functional specialization within the Fur family of metalloregulators. Biometals 20, 485–499. 10.1007/s10534-006-9070-7.

9. Oglesby-Sherrouse, A.G., and Murphy, E.R. (2013). Iron-responsive bacterial small RNAs: variations on a theme. Metallomics 5, 276–286. 10.1039/c3mt20224k.

10. Fillat, M.F. (2014). The FUR (ferric uptake regulator) superfamily: diversity and versatility of key transcriptional regulators. Arch Biochem Biophys 546, 41–52. 10.1016/j.abb.2014.01.029.

11. Hantke, K. (1981). Regulation of ferric iron transport in *Escherichia coli* K12: isolation of a constitutive mutant. Mol Gen Genet 182, 288–292. 10.1007/bf00269672.

12. Pandey, S.S., Patnana, P.K., Lomada, S.K., Tomar, A., and Chatterjee, S. (2016). Co-regulation of iron metabolism and virulence associated functions by iron and XibR, a novel iron binding transcription factor, in the plant pathogen *Xanthomonas*. PLoS Pathog 12, e1006019. 10.1371/journal.ppat.1006019.

13. Wu, S., Liu, J., Liu, C., Yang, A., and Qiao, J. (2020). Quorum sensing for population-level control of bacteria and potential therapeutic applications. Cell Mol Life Sci 77, 1319–1343. 10.1007/s00018-019-03326-8.

14. Wen, Y., Kim, I.H., Son, J.S., Lee, B.H., and Kim, K.S. (2012). Iron and quorum sensing coordinately regulate the expression of vulnibactin biosynthesis in *Vibrio vulnificus*. J Biol Chem 287, 26727–26739. 10.1074/jbc.M112.374165.

15. Chapalain, A., Vial, L., Laprade, N., Dekimpe, V., Perreault, J., and Déziel, E. (2013). Identification of quorum sensing-controlled genes in *Burkholderia ambifaria*. Microbiologyopen 2, 226–242. 10.1002/mbo3.67.

16. McRose, D.L., Baars, O., Seyedsayamdost, M.R., and Morel, F.M.M. (2018). Quorum sensing and iron regulate a two-for-one siderophore gene cluster in *Vibrio harveyi*. Proc Natl Acad Sci U S A 115, 7581–7586. 10.1073/pnas.1805791115.

17. Shah, N., Gislason, A.S., Becker, M., Belmonte, M.F., Fernando, W.G.D., and de Kievit, T.R. (2020). Investigation of the quorum-sensing regulon of the biocontrol bacterium *Pseudomonas chlororaphis* strain PA23. PLOS ONE 15, e0226232. 10.1371/journal.pone.0226232.

18. Lima-Bittencourt, C.I., Astolfi-Filho, S., Chartone-Souza, E., Santos, F.R., and Nascimento, A.M. (2007). Analysis of *Chromobacterium* sp. natural isolates from different Brazilian ecosystems. BMC Microbiol 7, 58. 10.1186/1471-2180-7-58.

19. Yang, C.H., and Li, Y.H. (2011). *Chromobacterium violaceum* infection: a clinical review of an important but neglected infection. J Chin Med Assoc 74, 435–441. 10.1016/j.jcma.2011.08.013.

20. Batista, J.H., and da Silva Neto, J.F. (2017). *Chromobacterium violaceum* pathogenicity: Updates and insights from genome sequencing of novel *Chromobacterium* species. Front Microbiol 8, 2213. 10.3389/fmicb.2017.02213.

21. Durán, N., and Menck, C.F. (2001). *Chromobacterium violaceum*: a review of pharmacological and industiral perspectives. Crit Rev Microbiol 27, 201–222. 10.1080/20014091096747.

22. Durán, N., Justo, G.Z., Durán, M., Brocchi, M., Cordi, L., Tasic, L., Castro, G.R., and Nakazato, G. (2016). Advances in *Chromobacterium violaceum* and properties of violacein-Its main secondary metabolite: A review. Biotechnol Adv 34, 1030–1045. 10.1016/j.biotechadv.2016.06.003.

23. Morohoshi, T., Kato, M., Fukamachi, K., Kato, N., and Ikeda, T. (2008). N-acylhomoserine lactone regulates violacein production in *Chromobacterium violaceum* type strain ATCC 12472. FEMS Microbiol Lett 279, 124–130. 10.1111/j.1574-6968.2007.01016.x.

24. McClean, K.H., Winson, M.K., Fish, L., Taylor, A., Chhabra, S.R., Camara, M., Daykin, M., Lamb, J.H., Swift, S., Bycroft, B.W., et al. (1997). Quorum sensing and *Chromobacterium violaceum*: exploitation of violacein production and inhibition for the detection of N-acylhomoserine lactones. Microbiology (Reading) 143 (*Pt 12*), 3703–3711. 10.1099/00221287-143-12-3703.

25. Stauff, D.L., and Bassler, B.L. (2011). Quorum sensing in *Chromobacterium violaceum*: DNA recognition and gene regulation by the CviR receptor. J Bacteriol 193, 3871–3878. 10.1128/jb.05125-11.

26. Mion, S., Carriot, N., Lopez, J., Plener, L., Ortalo-Magné, A., Chabrière, E., Culioli, G., and Daudé, D. (2021). Disrupting quorum sensing alters social interactions in *Chromobacterium violaceum*. NPJ Biofilms Microbiomes 7, 40. 10.1038/s41522-021-00211-w.

27. Evans, K.C., Benomar, S., Camuy-Vélez, L.A., Nasseri, E.B., Wang, X., Neuenswander, B., and Chandler, J.R. (2018). Quorum-sensing control of antibiotic resistance stabilizes cooperation in *Chromobacterium violaceum*. ISME J 12, 1263–1272. 10.1038/s41396-018-0047-7.

28. Batista, B.B., Santos, R., Ricci-Azevedo, R., and da Silva Neto, J.F. (2019). Production and uptake of distinct endogenous catecholate-type siderophores are required for iron acquisition and virulence in *Chromobacterium violaceum*. Infect Immun 87. 10.1128/iai.00577-19.

29. de Lima, V.M., Batista, B.B., and da Silva Neto, J.F. (2022). The regulatory protein ChuP connects heme and siderophore-mediated iron acquisition systems required for *Chromobacterium violaceum* virulence. Front Cell Infect Microbiol 12, 873536. 10.3389/fcimb.2022.873536.

30. Santos, R., Batista, B.B., and da Silva Neto, J.F. (2020). Ferric uptake regulator Fur coordinates siderophore production and defense against iron toxicity and oxidative stress and contributes to virulence in *Chromobacterium violaceum*. Appl Environ Microbiol 86. 10.1128/aem.01620-20.

31. Jacobs, M.A., Alwood, A., Thaipisuttikul, I., Spencer, D., Haugen, E., Ernst, S., Will, O., Kaul, R., Raymond, C., Levy, R., et al. (2003). Comprehensive transposon mutant library of *Pseudomonas aeruginosa*. Proc Natl Acad Sci U S A 100, 14339–14344. 10.1073/pnas.2036282100.

32. Lozano, G.L., Guan, C., Cao, Y., Borlee, B.R., Broderick, N.A., Stabb, E.V., and Handelsman, J. (2020). A Chemical Counterpunch: *Chromobacterium violaceum* ATCC 31532 produces violacein in response to translation-inhibiting antibiotics. mBio 11. 10.1128/mBio.00948-20.

33. Devescovi, G., Kojic, M., Covaceuszach, S., Cámara, M., Williams, P., Bertani, I., Subramoni, S., and Venturi, V. (2017). Negative regulation of violacein biosynthesis in *Chromobacterium violaceum*. Front Microbiol 8, 349. 10.3389/fmicb.2017.00349.

34. Swem, L.R., Swem, D.L., O’Loughlin, C.T., Gatmaitan, R., Zhao, B., Ulrich, S.M., and Bassler, B.L. (2009). A quorum-sensing antagonist targets both membrane-bound and cytoplasmic receptors and controls bacterial pathogenicity. Mol Cell 35, 143–153. 10.1016/j.molcel.2009.05.029.

35. Jones, C.M., Wells, R.M., Madduri, A.V., Renfrow, M.B., Ratledge, C., Moody, D.B., and Niederweis, M. (2014). Self-poisoning of *Mycobacterium tuberculosis* by interrupting siderophore recycling. Proc Natl Acad Sci U S A. 111, 1945–1950. 10.1073/pnas.1311402111.

36. Jin, Z., Li, J., Ni, L., Zhang, R., Xia, A., and Jin, F. (2018). Conditional privatization of a public siderophore enables *Pseudomonas aeruginosa* to resist cheater invasion. Nature Communications 9, 1383. 10.1038/s41467-018-03791-y.

37. Wang, L., Pan, Y., Yuan, Z.H., Zhang, H., Peng, B.Y., Wang, F.F., and Qian, W. (2016). Two-Component signaling system VgrRS directly senses extracytoplasmic and intracellular iron to control bacterial adaptation under iron depleted stress. PLoS Pathog 12, e1006133. 10.1371/journal.ppat.1006133.

38. Barragán, M.J., Blázquez, B., Zamarro, M.T., Mancheño, J.M., García, J.L., Díaz, E., and Carmona, M. (2005). BzdR, a repressor that controls the anaerobic catabolism of benzoate in *Azoarcus sp.* CIB, is the first member of a new subfamily of transcriptional regulators. J Biol Chem 280, 10683–10694. 10.1074/jbc.M412259200.

39. Durante-Rodríguez, G., Valderrama, J.A., Mancheño, J.M., Rivas, G., Alfonso, C., Arias-Palomo, E., Llorca, O., García, J.L., Díaz, E., and Carmona, M. (2010). Biochemical characterization of the transcriptional regulator BzdR from *Azoarcus sp*. CIB. J Biol Chem 285, 35694–35705. 10.1074/jbc.M110.143503.

40. Wang, H.C., Ko, T.P., Wu, M.L., Ku, S.C., Wu, H.J., and Wang, A.H. (2012). *Neisseria* conserved protein DMP19 is a DNA mimic protein that prevents DNA binding to a hypothetical nitrogen-response transcription factor. Nucleic Acids Res 40, 5718–5730. 10.1093/nar/gks177.

41. Alves, J.A., Leal, F.C., Previato-Mello, M., and da Silva Neto, J.F. (2022). A Quorum Sensing-Regulated Type VI Secretion System Containing Multiple Nonredundant VgrG Proteins Is Required for Interbacterial Competition in *Chromobacterium violaceum*. Microbiol Spectr 10, e0157622. 10.1128/spectrum.01576-22.

42. Mukherjee, S., Moustafa, D., Smith, C.D., Goldberg, J.B., and Bassler, B.L. (2017). The RhlR quorum-sensing receptor controls *Pseudomonas aeruginosa* pathogenesis and biofilm development independently of its canonical homoserine lactone autoinducer. PLoS Pathog 13, e1006504. 10.1371/journal.ppat.1006504.

43. Mukherjee, S., Moustafa, D.A., Stergioula, V., Smith, C.D., Goldberg, J.B., and Bassler, B.L. (2018). The PqsE and RhlR proteins are an autoinducer synthase-receptor pair that control virulence and biofilm development in *Pseudomonas aeruginosa*. Proc Natl Acad Sci U S A 115, E9411–e9418. 10.1073/pnas.1814023115.

44. Chernin, L. S., Winson, M. K., Thompson, J. M., Haran, S., Bycroft, B. W., Chet, I., Williams, P., and Stewart, G. S. (1998). Chitinolytic activity in *Chromobacterium violaceum*: substrate analysis and regulation by quorum sensing. J Bacteriol 180, 4435–4441. 10.1128/JB.180.17.4435-4441.1998.

45. Ross-Gillespie, A., Dumas, Z., and Kümmerli, R. (2015). Evolutionary dynamics of interlinked public goods traits: an experimental study of siderophore production in *Pseudomonas aeruginosa*. J Evol Biol 28, 29–39. 10.1111/jeb.12559.

46. Popat, R., Harrison, F., da Silva, A.C., Easton, S.A., McNally, L., Williams, P., and Diggle, S.P. (2017). Environmental modification via a quorum sensing molecule influences the social landscape of siderophore production. Proc Biol Sci 284. 10.1098/rspb.2017.0200.

47. Eickhoff, M.J., and Bassler, B.L. (2020). *Vibrio fischeri* siderophore production drives competitive exclusion during dual-species growth. Mol Microbiol 114, 244–261. 10.1111/mmi.14509.

48. Zhang, Y., Gao, J., Wang, L., Liu, S., Bai, Z., Zhuang, X., and Zhuang, G. (2018). Environmental Adaptability and Quorum Sensing: Iron uptake regulation during biofilm formation by *Paracoccus denitrificans*. Appl Environ Microbiol 84. 10.1128/aem.00865-18.

49. Lin, J.S., and Lai, E.M. (2017). Protein-Protein Interactions: Yeast Two-Hybrid System. Methods Mol Biol 1615, 177–187. 10.1007/978-1-4939-7033-9_14.

50. da Silva Neto, J.F., Negretto, C.C., and Netto, L.E. (2012). Analysis of the organic hydroperoxide response of *Chromobacterium violaceum* reveals that OhrR is a cys-based redox sensor regulated by thioredoxin. PLoS One 7, e47090. 10.1371/journal.pone.0047090.

51. Schwyn, B., and Neilands, J.B. (1987). Universal chemical assay for the detection and determination of siderophores. Anal Biochem 160, 47–56. 10.1016/0003-2697(87)90612-9.

52. Pandey, A., and Sonti, R.V. (2010). Role of the FeoB protein and siderophore in promoting virulence of *Xanthomonas oryzae pv. oryzae* on rice. J Bacteriol 192, 3187–3203. 10.1128/jb.01558-09.

53. Ten-Caten, F., Vêncio, R.Z.N., Lorenzetti, A.P.R., Zaramela, L.S., Santana, A.C., and Koide, T. (2018). Internal RNAs overlapping coding sequences can drive the production of alternative proteins in archaea. RNA Biol 15, 1119–1132. 10.1080/15476286.2018.1509661.

54. De Souza, W., Carvalho, B.D.S., Lopes-Cendes, I. (2018). Rqc: A Bioconductor Package for Quality Control of High-Throughput Sequencing Data. Journal of Statistical Software 87, Code Snippet 2. 10.18637/jss.v087.c02.

55. Bolger, A.M., Lohse, M., and Usadel, B. (2014). Trimmomatic: a flexible trimmer for Illumina sequence data. Bioinformatics 30, 2114–2120. 10.1093/bioinformatics/btu170.

56. Kim, D., Paggi, J.M., Park, C., Bennett, C., and Salzberg, S.L. (2019). Graph-based genome alignment and genotyping with HISAT2 and HISAT-genotype. Nat Biotechnol 37, 907–915. 10.1038/s41587-019-0201-4.

57. de Araújo, H.L., Martins, B.P., Vicente, A.M., Lorenzetti, A.P.R., Koide, T., and Marques, M.V. (2021). Cold regulation of genes encoding ion transport systems in the oligotrophic bacterium *Caulobacter crescentus*. Microbiol Spectr 9, e0071021. 10.1128/Spectrum.00710-21.

58. Lawrence, M., Huber, W., Pagès, H., Aboyoun, P., Carlson, M., Gentleman, R., Morgan, M.T., and Carey, V.J. (2013). Software for computing and annotating genomic ranges. PLoS Comput Biol 9, e1003118. 10.1371/journal.pcbi.1003118.

59. Love, M.I., Huber, W., and Anders, S. (2014). Moderated estimation of fold change and dispersion for RNA-seq data with DESeq2. Genome Biol 15, 550. 10.1186/s13059-014-0550-8.

60. Livak, K.J., Schmittgen, T.D. (2001) Analysis of relative gene expression data using real-time quantitative PCR and the 2(-Delta Delta C(T)) Method. Methods 25, 402–408. 10.1006/meth.2001.1262.

